# Ablation of *Max* expression induces meiotic onset in sexually undifferentiated germ cells

**DOI:** 10.1101/2023.06.10.544477

**Authors:** Ayumu Suzuki, Kousuke Uranishi, Masazumi Nishimoto, Yosuke Mizuno, Seiya Mizuno, Satoru Takahashi, Robert N. Eisenman, Akihiko Okuda

## Abstract

Meiosis is a specialized type of cell division that occurs only in germ cells physiologically. We have previously demonstrated that MYC-associated factor X (MAX) is involved in blocking ectopic and precocious meiotic onsets in embryonic and germline stem cells, respectively, as a central component of the PRC1 subtype PRC1.6. In this study, we investigated the role of the *Max* gene in germ cells *in vivo*. Our data revealed that mitotically active germ cell-specific disruption of *Max* was coupled to meiotic onset in male and female germ cells. However, such *Max*-null germ cells did not undergo meiotic processes progressively, but were stalled at its relatively early stages and eventually eliminated by apoptosis. Our data also revealed that *Max*, which is generally known as an obligate heterodimerization partner for MYC and MXD transcription factors, showed high expression in sexually undifferentiated male and female germ cells, but female germ cells exhibited an abrupt decline in its expression at the timing of or immediately prior to physiological meiotic onset. Moreover, computational analyses identified the regulatory region that supported high levels of *Max* expression in sexually undifferentiated germ cells.

## Introduction

Germ cells are the only cell type capable of transmitting genetic information to the next generation. Meiosis as a fundamental process of gametogenesis is a specialized type of cell division that halves the number of chromosomes (Handel and Schimenti 2010; Kimble 2011; Suzuki et al. 2017; Spiller and Bowles 2022). While the molecular bases of most events that occur during meiosis are highly conserved among eukaryotes, the regulatory mechanisms that control meiotic onset in germ cells in mammals are largely unknown because of their limited evolutionary conservation (Marston and Amon 2004; Kimble 2011; Lesch and Page 2012; Farini and De Felici 2022; Spiller and Bowles 2022). Dysregulation of meiotic genes in germ cells is closely linked to infertility and cancer (Carroll et al. 2018; Feichtinger and McFarlane 2019; Nicholls and Page 2021; Farini and De Felici 2022; Kitayama et al. 2022; Lingg et al. 2022), underscoring the importance of elucidating the detailed molecular mechanisms that control meiotic onset in mammals.

During fetal development, primordial germ cells (PGCs) are generated from several epiblast cells located in the posterior/proximal corner at approximately embryonic day (E) 6.25 and migrate through the hindgut to the genital ridge by approximately E10.5, remaining sexually undifferentiated until approximately E11.5 [reviewed in (Spiller and Bowles 2022)]. Whereas female germ cells undergo meiosis at approximately E13.5, male germ cells do not undergo meiosis during the embryonic stage and onset occurs after birth. The *Signaled by retinoic acid 8* (*Stra8*) gene is a crucial regulator of meiotic onset in male and female germ cells (Baltus et al. 2006; Bowles et al. 2006; Koubova et al. 2006; Anderson et al. 2008). Although the molecular mechanism by which STRA8 for sustains early stages of meiotic processes have long been enigmatic, identification of MEIOSIN as a binding partner of STRA8 revealed that STRA8, by interacting with MEIOSIN, functions as a transcriptional activator that activates numerous meiosis-related genes including *Stra8* and *Meiosin* genes themselves (Ishiguro et al. 2020). However, it remains unclear as to whether epigenetic regulation allows the STRA8/MEIOSIN transcriptional complex to perform its function effectively.

Although MYC-associated factor X (MAX) is generally known as an obligate partner of MYC to activate a wide range of genes involved in cell proliferation and metabolism (Eilers and Eisenman 2008; Carroll et al. 2018), MAX also heterodimerizes with the bHLHZ region of MGA and the resultant MGA-MAX heterodimer is a critical component of polycomb repressive complex (PRC) 1.6, a subtype of PRC1 that regulates the transcriptionally repressed chromatin state (Gao et al. 2012). Although the physiological significance is unknown, we have previously demonstrated that mouse embryonic stem cells (ESCs) have the potential for meiosis onset, and PRC1.6 prevents ectopic onset of meiosis in ESCs (Suzuki et al. 2016). Indeed, our data demonstrated that knockout of the *Max* gene in ESCs was accompanied by cytological changes reminiscent of germ cells undergoing meiotic prophase I due to potent derepression of many meiosis-related genes such as *Sycp3* and *Dazl*. Furthermore, we and others have previously demonstrated that PRC1.6 binds to the promoters of meiosis-related genes and represses their expression in ESCs (Kehoe et al. 2008; Hisada et al. 2012; Qin et al. 2012; Leseva et al. 2013; Maeda et al. 2013; Suzuki et al. 2016; Yang et al. 2016; Endoh et al. 2017; Huang et al. 2018; Stielow et al. 2018; Uranishi et al. 2021). Moreover, we have demonstrated that functional impairment of PRC1.6 in germline stem cells (GSCs) induces meiosis-like phenotypic changes (Suzuki et al. 2016). Additionally, a report has demonstrated a direct link between meiotic onset and PRC1, in which knockout of the gene encoding RING1B, a common component of all PRC1 subtypes, provokes precocious meiotic entry in female primordial germ cells (PGCs) (Yokobayashi et al. 2013). Considering these findings, we hypothesized that PRC1.6 (containing MAX and MGA) normally acts to restrict meiotic entry in germ cells.

To test this hypothesis, we conducted PGC-specific knockout of *Max* and examined the consequences. We found that sexually undifferentiated germ cells showed meiotic entry upon *Max* gene disruption, but such *Max*-null germ cells were stalled at relatively early stages of meiosis and eventually eliminated by apoptosis. We also found that *Max* had relatively higher expression in mitotically active male and female germ cells physiologically compared with somatic cells surrounding the germ cells. Subsequently, female, but not male, germ cells showed an abrupt decline in *Max* expression at the time of, or immediately prior to, physiological meiotic onset that occurs specifically in female germ cells at midgestation stage to mitigate PRC1.6 inhibition of meiosis. Moreover, we identified the regulatory region of *Max* that contributes to its characteristic expression profile.

## Results

### Meiotic onset is preceded by an abrupt decline in *Max* expression in female PGCs

To explore the relationship between *Max* expression and meiotic onset in germ cells *in vivo*, we assessed Max protein levels by immunohistochemistry in male and female germ cells at E11.5 that expressed pluripotency marker Oct4 and were therefore not sexually differentiated (Fig. 1A). Oct4-positive male and female germ cells at this stage had significantly strong Max protein staining compared with surrounding Oct4-negative somatic cells. However, when examined at E13.5, Max protein staining in meiotic female germ cells at preleptotene and early leptotene stages judged by the SYCP3 staining pattern was equivalent and even weaker compared with that in surrounding somatic cells, respectively (Fig. 1B). Quantification of Max protein levels in germ cells during the midgestation stage revealed that female germ cells showed an abrupt decline between E11.5 and E13.5 that led to levels lower than those in somatic cells, contrasting the continuous gradual decline in male germ cells that do not undergo meiosis during this stage (Fig. 1C). Analyses using publicly available transcriptome analysis data (Sabour et al. 2011) also revealed significantly different *Max* expression profiles in male and female PGCs. The *Max* expression level started an abrupt decline in female PGCs at E11.5, leading to less than half the level by meiotic onset at E13.5, which increased from E16.5. However, male PGCs showed a gradual decrease in the *Max* mRNA level in a constant manner during these periods (Supplemental Fig. S1A). These data indicated that the abrupt and gradual declines in the *Max* mRNA level in female and male PGCs, respectively, which occurred between E11.5 and E13.5, were largely reflected by their Max protein levels.

**Figure 1.**
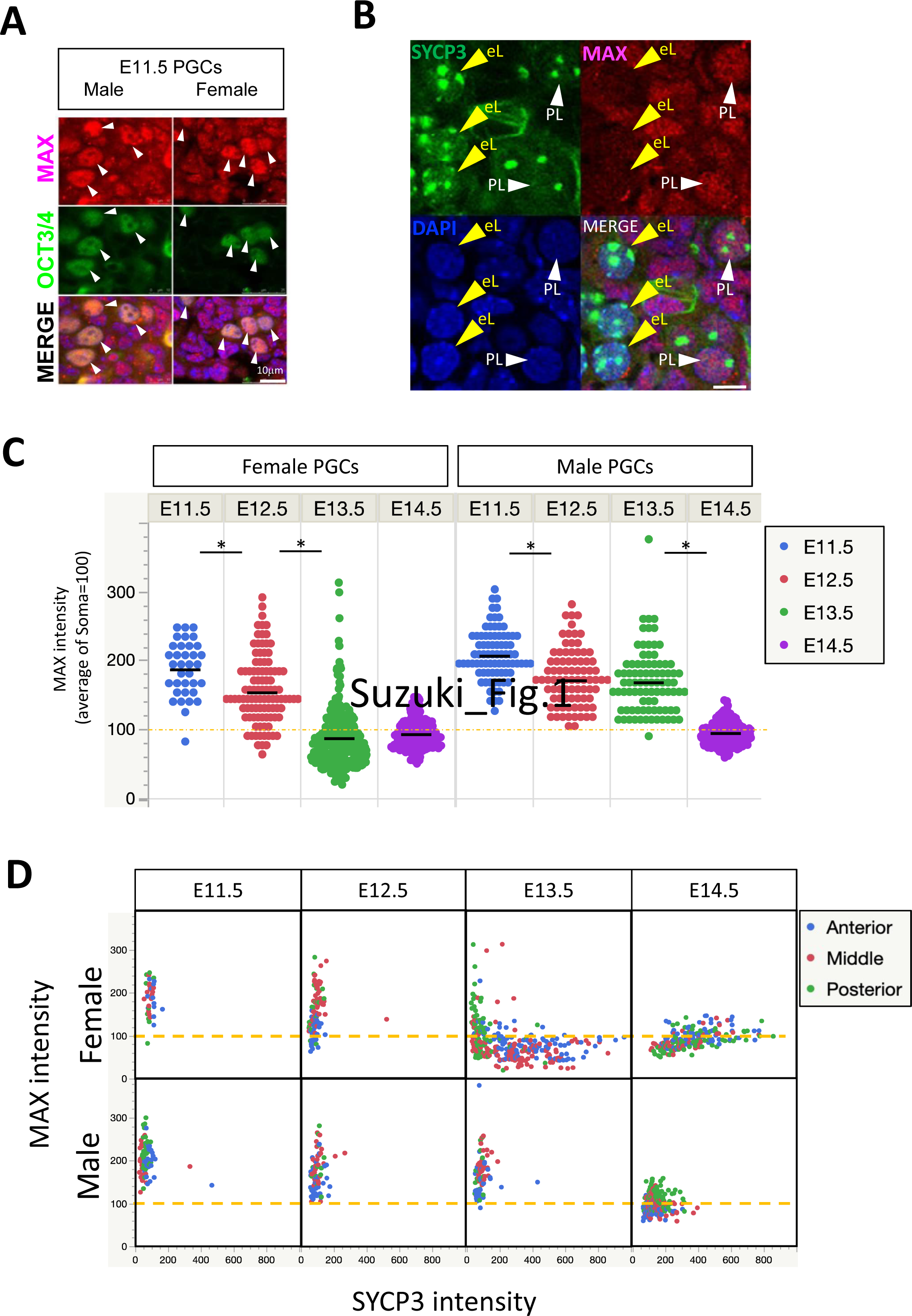
Spatiotemporal alterations of Max protein levels in premeiotic male and female germ cells. (A) Double immunofluorescence staining of MAX and OCT4 (POU5F1) in male and female gonads at E11.5. Arrowheads indicate each OCT4-positive germ cell. (B) Double immunofluorescence analyses. Gonadal sections of female germ cells at E13.5 were immunostained for MAX and SYCP3, and counterstained with DAPI. PL: preleptotene; eL: early leptotene. (C) Plots of the intensity of individual Max staining signals in male and female germ cells. The relative intensity of Max staining in individual germ cells in male and female gonads at E11.5, E12.5, E13.5, and E14.5 was determined using ImageJ software. The mean Max signal intensity in somatic cells was arbitrarily set to 100. (D) Scatter plots representing the intensity of Max and Sycp3 staining signals in individual male and female germ cells at E13.5 localized at the anterior (blue dot), middle (red dot), or posterior (green dot) portion of gonads. The intensity of Max and Sycp3 staining signals in somatic cells was set to 100.

Meiosis occurs in female gonads along the anterior–posterior axis. It occurs restrictively in germ cells located at the anterior portion of the gonad at approximately E13.5, and then the region with germ cells undergoing meiosis expands and/or moves to the posterior region in a wave-like fashion (Menke et al. 2003; Yao et al. 2003; Bullejos and Koopman 2004). Therefore, we used publicly available global gene expression data (Soh et al. 2015) to examine whether expression levels of the *Max* gene as well as pluripotency- and meiosis-related genes were influenced by the location of female germ cells in the gonad. Female germ cells located at the posterior portion of the gonad at E13.5 had relatively higher expression levels of *Max* and pluripotency genes (*Oct4* and *Nanog)*, but lower expression levels of meiosis-related genes (*Dazl*, *Stra8*, and *Sycp3*) compared with germ cells located at the anterior portion (Supplemental Fig. S1B). Next, we examined MAX and SYCP3 protein levels in female germ cells by double immunohistochemistry in relation to the spatiotemporal regulation of meiotic entry. Female germ cells in the posterior portion of the gonad also had a delayed reduction and increase in Max and Sycp3 protein levels, respectively, compared with those in the anterior portion (Fig. 1D), strengthening the notion that the Max protein decline and meiosis were closely linked events. However, such an inverse correlation between them was no longer evident at E14.5, but they appeared to become positively correlated. In contrast to female germ cells, male germ cells, which do not produce Sycp3 protein at appreciable levels during embryonic stages, showed a continuous decrease in the Max protein level between E11.5 and E14.5, but there was no spatiotemporal difference. Immunocytochemical analyses of female gonadal cells dissociated to single cells revealed the lowest Max protein level in the early leptotene stage. However, the Max protein level increased during subsequent leptotene and zygotene stages (Supplemental Fig. S2), which was in line with *Max* expression starting to increase in female germ cells from the midgestation stage (Supplemental Fig. S1A).

### *Max* gene disruption is coupled to meiotic onset in sexually undifferentiated male and female PGCs

To further explore the relationship between *Max* expression and meiotic onset in germ cells, we conducted germ cell-specific knockout of the *Max* gene. To this end, we generated mice with a *Max* gene that is subject to disruption specifically in sexually undifferentiated germ cells by tamoxifen administration. These mice harbor a homozygous *Max*-floxed allele (*Max*^fl/fl^ mice) (Mathsyaraja et al. 2019) and hemizygous Oct4dPE-CreERT2 transgene expressed under the control of a distal enhancer of the *Oct4* gene (Fig. 2A). We examined alterations in the expression levels of meiosis-related genes (*Sycp3* and *Meiosin*) due to *Max* gene disruption in male and female germ cells by tamoxifen administration (Fig. 2B). Significant activation of both genes in male germ cells by induction of *Max* gene disruption via tamoxifen administration was evident between E10.5 and E14.5. Such activation was most prominent at E12.5, but the magnitude of activation became less significant at E13.5 and E14.5. Female germ cells also showed similar activation of these genes after induction of *Max* gene disruption. However, such activation was observed only before sexual differentiation (E11.5 and E12.5). Indeed, expression of these genes was comparable and lower in female germ cells subjected to Cre-mediated *Max* gene disruption compared with controls without CreERT2 cDNA at E13.5 and E14.5, respectively (Fig. 2B). These data indicated that forced meiotic onset that were induced in female germ cells precociously did not act in concert with physiological meiosis initiating at approximately E13.5, but instead acted to limit the progression of physiological meiotic processes. Immunohistochemical analyses showed a high level of MAX, but not SYCP3, at E11.5 in nuclei of germ cells of male and female gonads in control mice. However, tamoxifen administration-mediated *Max* gene disruption gave rise to germ cells with no Max protein signal (Fig. 2C, D). More importantly, such Max signal loss was in most cases accompanied by acquisition of a significant Sycp3 signal in male germ cells and even stronger Sycp3 signal in female germ cells at E11.5 and E12.5 (Fig. 2C, D, Fig. S3), during which male and female germ cells are not sexually differentiated. However, such a significant Sycp3 protein signal was hardly detected in *Max* expression-ablated male PGCs at E13.5 and E14.5 (Fig. 2C, D, Supplemental Fig. S3). These results indicated that *Max* expression-ablated male germ cells did not maintain the meiosis-like cytological changes observed before sexual differentiation because of the reduction in expression of meiosis-related genes during meiotic processes. Unlike male PGCs, completely Max protein signal-null female PGCs retained the Sycp3 signal even at E13.5 and E14.5. However, the immunostaining pattern of SYCP3 revealed that germ cells at relatively earlier (preleptotene and leptotene) and advanced (zygotene and pachytene) stages were respectively enriched and decreased among germ cells subjected to *Max* gene disruption compared with control female germ cells with physiological meiosis onset (Fig. 2E). Taken together, complete ablation of *Max* expression was sufficient to induce meiotic onset forcedly in sexually undifferentiated male and female PGCs. However, such *Max*-null germ cells failed to undergo subsequent meiotic processes progressively and were stalled immediately after forced induction of meiotic onset. Next, we examined the effect of ablating *Max* expression on migrating PGCs. The Sycp3 signal intensity was extremely low in most Max protein signal-null migrating germ cells at E10.5 (Supplemental Fig. S4A). However, some completely *Max* expression-ablated germ cells that had migrated to the vicinity of the genital ridge showed a relatively higher Sycp3 protein signal (Supplemental Fig. S4). These results indicated that germ cells had acquired competence to respond to forced meiotic induction by *Max* expression ablation during their migration.

**Figure 2.**
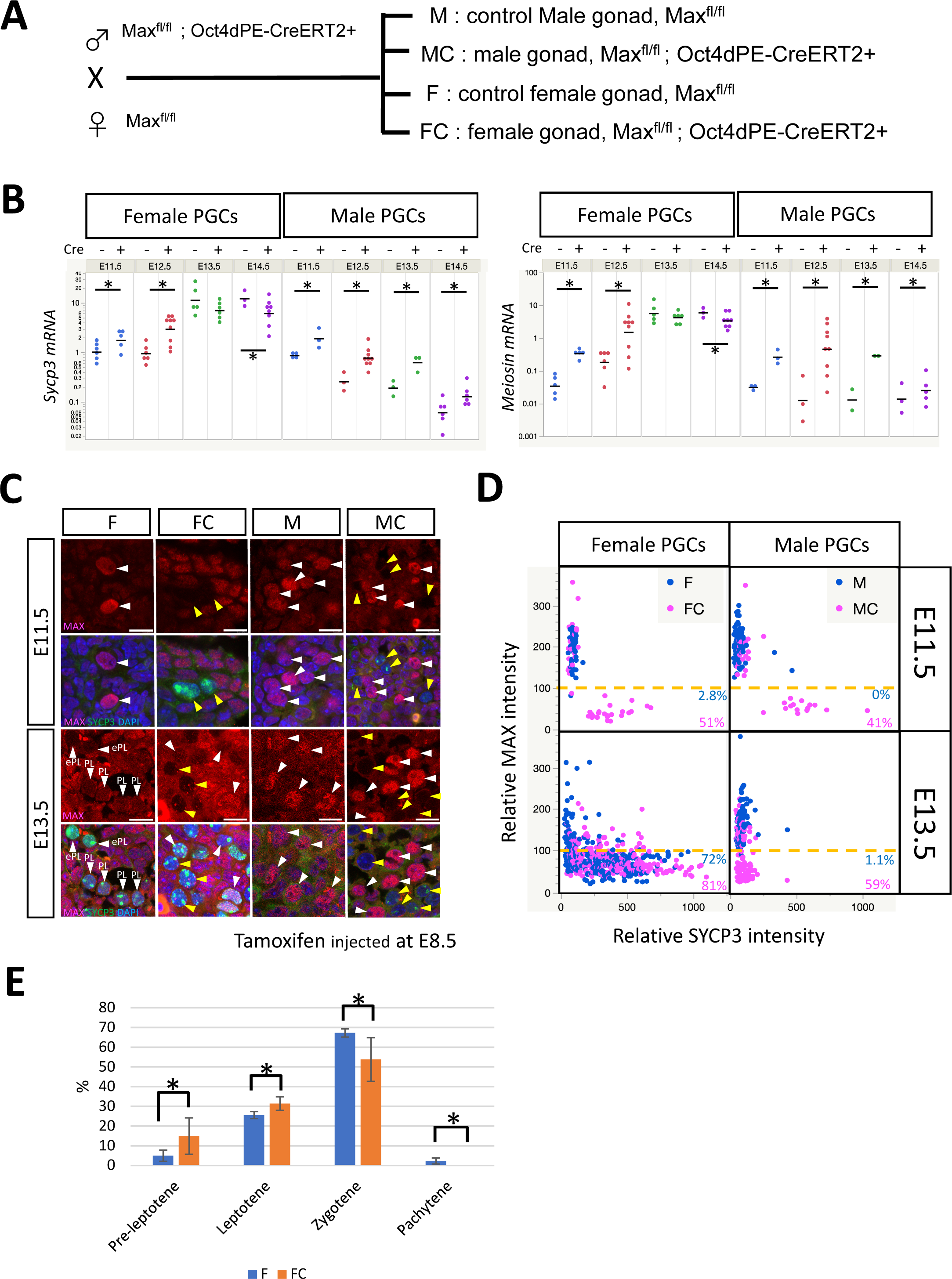
*Max* conditional knockout in sexually undifferentiated premeiotic germ cells induces meiotic onset. (A) Mating scheme to generate *Max* conditional knockout mice. Max^fl/fl^;Oct4dPE-CreERT2^+^ male mice were crossed with Max^fl/fl^ female mice. Male and female embryos with CreERT2 cDNA were designated as MC and FC, while male and female embryos without the cDNA were designated as M and F, respectively. (B) Examination of the effect of germ cell-specific disruption of the *Max* gene on expression of meiosis-related genes. *Sycp3* and *Meiosin* mRNA levels in germ cells from M, MC, F, and FC embryos at E10.5, E11.5, E12.5, E13.5, and E14.5 were quantified by quantitative PCR. Data represent the mean ± standard deviation of at least three independent experiments. \**P* < 0.05, Student’s t-test. (C) Double immunostaining of germ cells subjected to Cre-mediated *Max* gene disruption by tamoxifen administration and their control counterparts without CreERT2 cDNA. Fetal female and male embryos with or without CreERT2 cDNA were treated with tamoxifen at E8.5 and then genital ridges were isolated from embryos at either E11.5 or E13.5. Genital ridges were sectioned and subjected to immunostaining of MAX (magenta) and SYCP3 (green), and counterstained with DAPI (Blue). Each germ cell is indicated by an arrowhead. Yellow arrowheads indicate germ cells with complete elimination of the Max protein signal by tamoxifen administration. g, PL, and eL indicate sexually undifferentiated germ cells, preleptotene, and early leptotene, respectively. (D) Scatter plot representing the intensity of Max and Sycp3 staining signals in individual male and female germ cells at E11.5 and E13.5. The proportions of germ cells with a lower Max protein signal, but higher Sycp3 protein signal compared with somatic cells are shown. (E) Proportions of germ cells in meiotic prophase l stages in gonads from F (n=3, 305 total cells) and FC (n=4, 456 total cells) embryos at E14.5. Germ cells were immunostained for SYCP3 after spreading onto a glass slide and classified into four stages (preleptotene, leptotene, zygotene, and pachytene) by the SYCP3 staining pattern. The difference in the proportion between F and FC mice at each stage was examined statistically by Student’s t-test. \**P* < 0.05.

To eliminate the possibility that forced meiotic onset in sexually undifferentiated *Max*-null male and female germ cells did not represent a primary, but secondary consequence of certain abnormalities in sexual differentiation, such as precocious sexual differentiation and sex reversal, we examined whether *Max* expression ablation altered the expression of genes related to sexual differentiation. Quantitative PCR analyses revealed that *Sox9*, which is crucially involved in male-specific characteristics (Kent et al. 1996; Sekido and Lovell-Badge 2008), did not have noticeably altered expression (Fig. S5A). Similarly, *Foxl2*, which plays indispensable roles in oocyte development and meiotic progression during the embryonic stage (Ottolenghi et al. 2005; Uhlenhaut et al. 2009; Tucker 2022), did not have appreciably altered expression (Fig. S5B). These data indicated that *Max* gene disruption in germ cells induced meiosis forcedly without affecting the expression of sexual differentiation-related genes in surrounding somatic cells.

### Global expression analyses of the effects of *Max* gene disruption on embryonic germ cells

To examine alterations in the gene expression profile caused by embryonic germ cell-specific *Max* gene disruption, we conducted DNA microarray analyses to compare global expression profiles between germ cells subjected to CRE recombinase-mediated *Max* gene disruption and their control counterparts without CreERT2 cDNA (Fig. 3A). Although genes upregulated by *Max* gene disruption in female PGCs did not substantially overlap with those in male PGCs (Fig. 3B), gene ontology (GO) analyses revealed over-representation of genes involved in meiosis in all three gene sets, i.e., genes commonly activated in male and female PGCs and those activated only in either male or female PGCs (Fig. 3C). It was noteworthy that substantial overlap was evident between these genes and genes denoted as germline reprogramming-responsive genes whose transcriptional activation is closely linked to extensive epigenetic reprogramming in germ cells at E10.5–E11.5 via DNA demethylation of their promoters with high CpG content (Fig. 3D) (Mochizuki et al. 2021). Although MAX was originally cloned as an obligate heterodimerization partner of Myc proteins that are predominantly associated with transcriptional activation of genes involved in cell proliferation and metabolic processes (Eilers and Eisenman 2008; Carroll et al. 2018), the vast majority of genes defined as cell type-independent targets of the Myc/Max transcriptional complex (Ji et al. 2011) did not show significantly altered expression (Supplemental Fig. S6). These results indicated that the major role of MAX at these stages was not to promote cell proliferation and cellular metabolism as an interacting partner of the Myc transcription factor.

**Figure 3.**
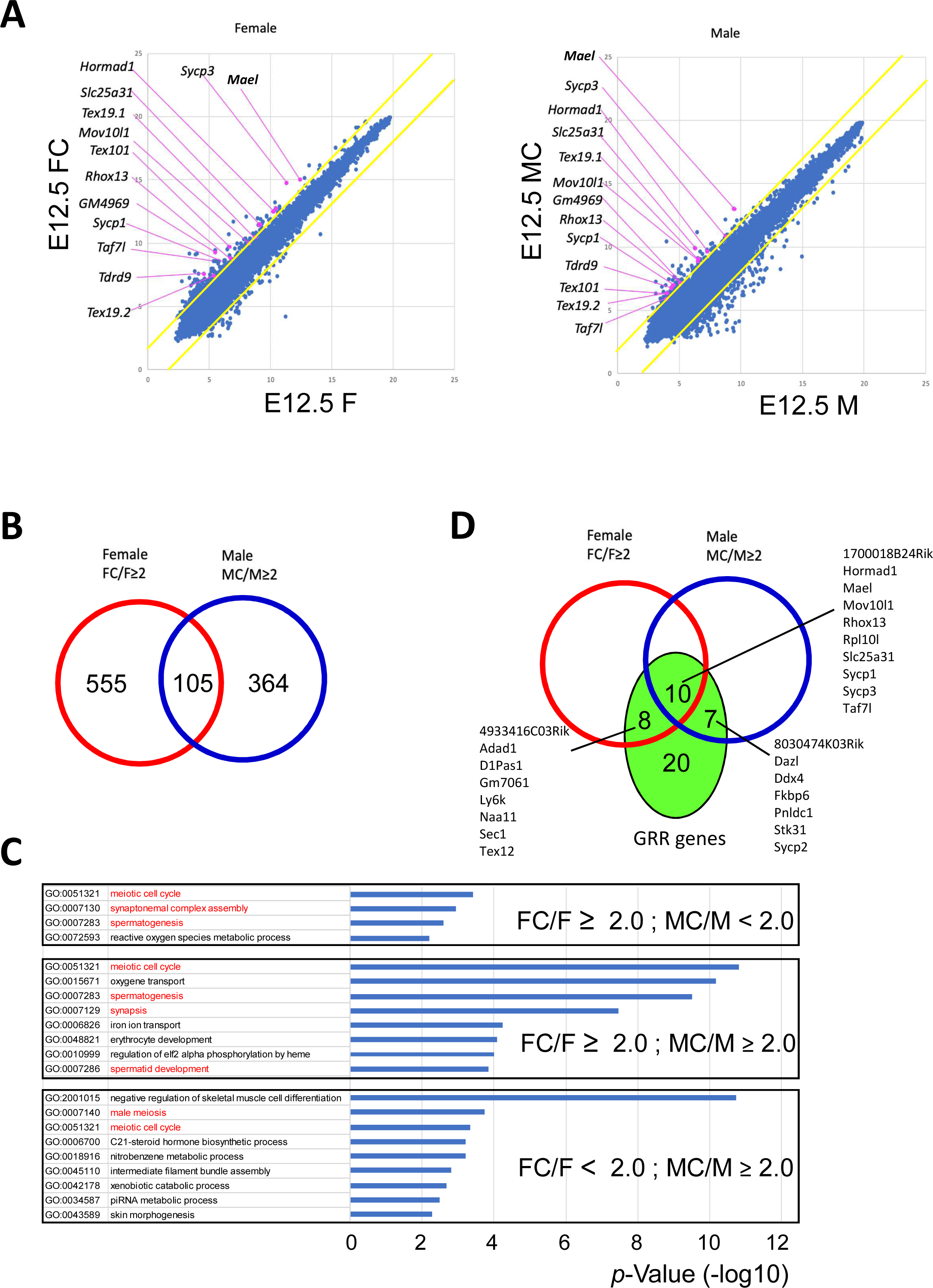
Alterations in the global expression profile of germ cells due to *Max* gene disruption. (A) Scatter plot of DNA microarray data from germ cells subjected to Cre-mediated *Max* gene knockout and their control counterparts without CreERT2 cDNA. Left and right panels show data from female and male germ cells at E13.5, respectively. Numerical values along X- and Y-axes are the log_2_-transformed signal intensity of the microarray data. Upper and lower yellow diagonal lines in each panel indicate the boundaries of 2-fold up- and down-regulation due to *Max* gene disruption, respectively. Meiosis-related genes upregulated by *Max* gene disruption are individually marked by pink dots with gene symbols. (B) Venn diagram comparing genes suggested to be activated by >2-fold after Cre-mediated *Max* gene disruption in female and male germ cells at E12.5. (C) GO analyses of activated genes upregulated by >2-fold after Cre-mediated *Max* gene disruption. Upregulated genes by >2-fold commonly in both male and female germ cells (middle panel) and those specifically in either male (bottom panel) or female (upper panel) germ cells were respectively subjected to GO analyses. Terms related to gametogenesis are indicated with red letters. (D) Venn diagram showing significant overlap between genes activated by *Max* gene disruption in germ cells and genes designated as germline reprogramming-responsive genes (Mochizuki et al. 2021).

### Germ cells with meiosis onset induced by *Max* expression ablation are eventually eliminated by apoptosis

Because male and female germ cells with meiosis onset induced by forced *Max* expression ablation were stalled in early stages of meiosis, we elucidated the subsequent fate of such meiotically arrested germ cells. Although total germ cell numbers in genital ridges assessed as TRA98-positive cells were unaffected by tamoxifen administration-mediated *Max* gene disruption at E12.5 in male and female embryos, a significant reduction in the germ cell number became evident in both sexes at E14.5 (Fig. 4A, B). To examine whether this reduction was due to apoptosis of *Max* expression-ablated germ cells, we immunostained cleaved PARP-1, which is an excellent apoptotic marker in germ cells (Anzar et al. 2002; Nguyen et al. 2020). In accordance with our hypothesis, there were significantly more cleaved PARP-1-positive germ cells at E14.5 among germ cells subjected to CRE recombination-mediated *Max* gene disruption compared with controls without CreERT2 cDNA in both male and female cases (Fig. 4C, Supplemental Fig. S7). Taken together, our data indicated that male and female germ cells that underwent ectopic and precocious meiosis, respectively, induced by *Max* expression ablation were stalled at early stages in their meiotic processes, and such germ cells were eventually eliminated by apoptotic cell death. Consistent with this notion, an obviously skewed distribution towards relatively earlier stages of meiotic processes among germ cells subjected to *Max* gene disruption compared with control germ cells at E14.5 (Fig. 2E) was no longer evident at E18.5 (Fig. 4D).

**Figure 4.**
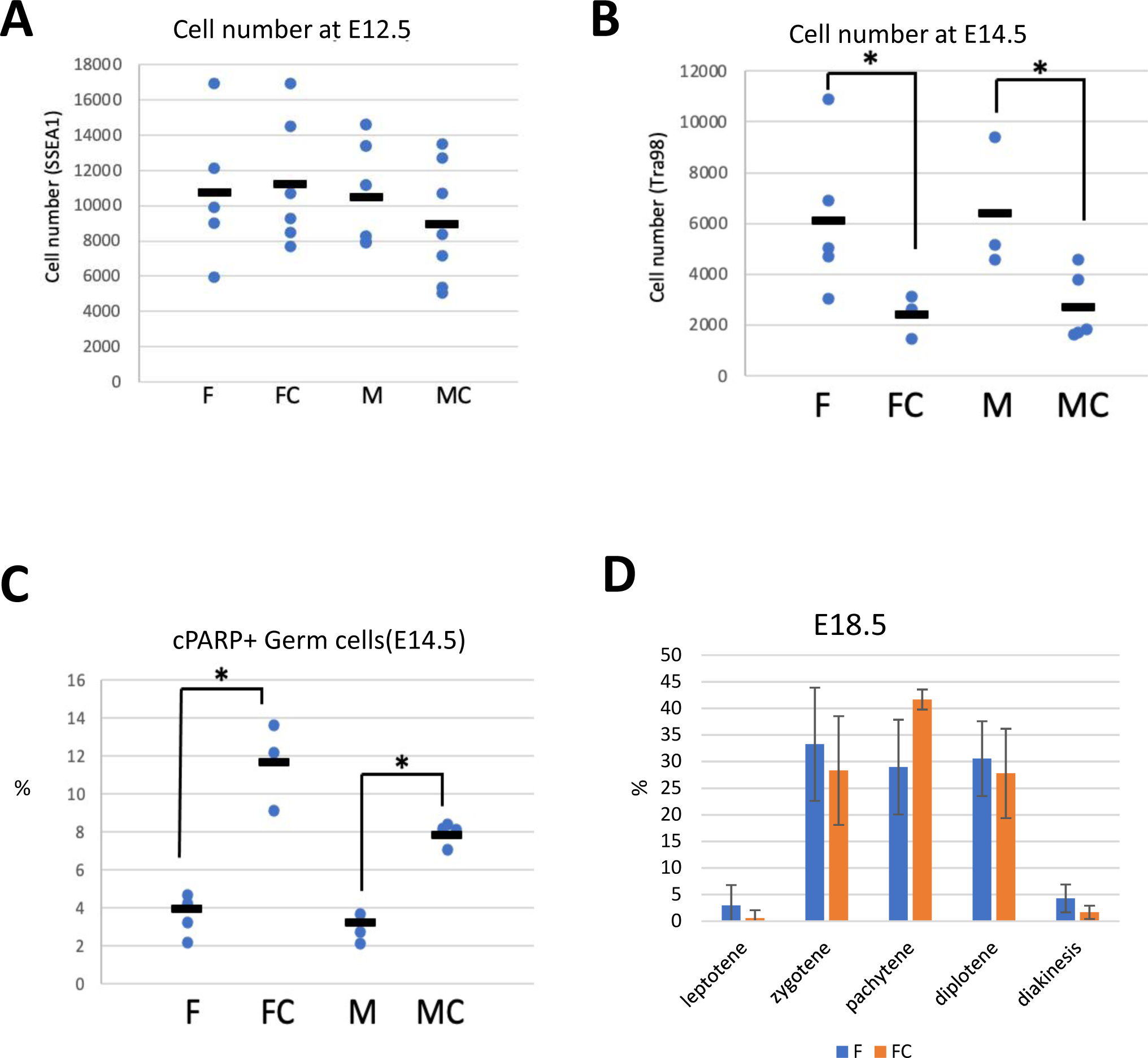
Apoptotic phenotype of primordial germ cells undergoing meiosis forcedly by *Max* expression ablation. (A, B) Total germ cell numbers assessed by TRA98 staining in gonads of M, MC, F, and FC mouse embryos at E12.5 (A) and E14.5 (B). (C) Percentages of cPARP-positive germ cells in gonads of M, MC, F, and FC mice at E14.5. (D) Proportions of germ cells in meiotic prophase l stages in E18.5 gonads of F and FC embryos. Germ cells were immunostained for SYCP3 after spreading onto a glass slide and their meiotic stages were classified as described in Fig. 2E.

### Identification of the regulatory region that supports high expression of *Max* in pluripotent embryonic stem cells and pre-meiotic germ cells

Similar to the *Max* gene, several pluripotency marker genes that encode transcription factors, such as *Oct4* and *Nanog*, are expressed in pre-meiotic germ cells and ESCs. Therefore, we hypothesized that *Max* contains a germ cell/ESC cell-type specific regulatory element that functions by recruiting commonly expressed pluripotency factors. To test this hypothesis, we searched for genomic regions localized in the vicinity of *Max* that were bound by certain pluripotency factors in mouse ESCs using ChIP-Atlas, a database of ChIP-seq data for meta-analysis. As a result, we identified a genomic region that met these criteria (Fig. 5A). This genomic region, which was named MUR (*<u>M</u>ax* <u>U</u>pstream <u>R</u>egion) was located at approximately 3 kb upstream of the transcription start site of *Max*. Analyses using publicly available DNase-seq data (Li et al. 2018) confirmed that MUR had an open chromatin configuration in ESCs and germ cells, but not in somatic cells (Supplemental Fig. S8A). These data supported the hypothesis that MUR was involved in transcriptional activation of ESCs and pre-meiotic germ cells. To evaluate transcription-stimulating activity of MUR in ESCs, we conducted luciferase assays using a luciferase reporter carrying MUR. The reporter with the MUR sequence induced 2- or 3-fold higher luciferase activity than the reporter without MUR in ESCs, whereas no statistical difference in luciferase activity due to the presence of MUR was evident in mouse embryonic fibroblasts (Fig. 5B). Next, we generated mice lacking MUR in their genome (dMUR) to assess the function of MUR to support *Max* expression in pre-meiotic germ cells. Although this deletion did not affect the germ cell number (Supplemental Fig. S8B), immunohistochemical analyses of MAX revealed that, compared to wild-type male and female germ cells, sexually undifferentiated germ cell mutants without MUR at E11.5 showed a significantly lower Max protein signal that was comparable with that observed in surrounding somatic cells (Fig. 5C). This observation indicated that MUR contributed to the high *Max* expression in ESCs and pre-meiotic germ cells. Quantification of *Max* mRNA supported this notion (Fig. 5D). However, unlike complete disruption of the *Max* gene, the decline in *Max* expression induced by MUR deletion was not accompanied by induction of meiotic gene expression (Fig. 5E). These data indicated that the decline in *Max* expression comparable with that in somatic cells was not sufficient to induce meiosis forcedly.

**Figure 5.**
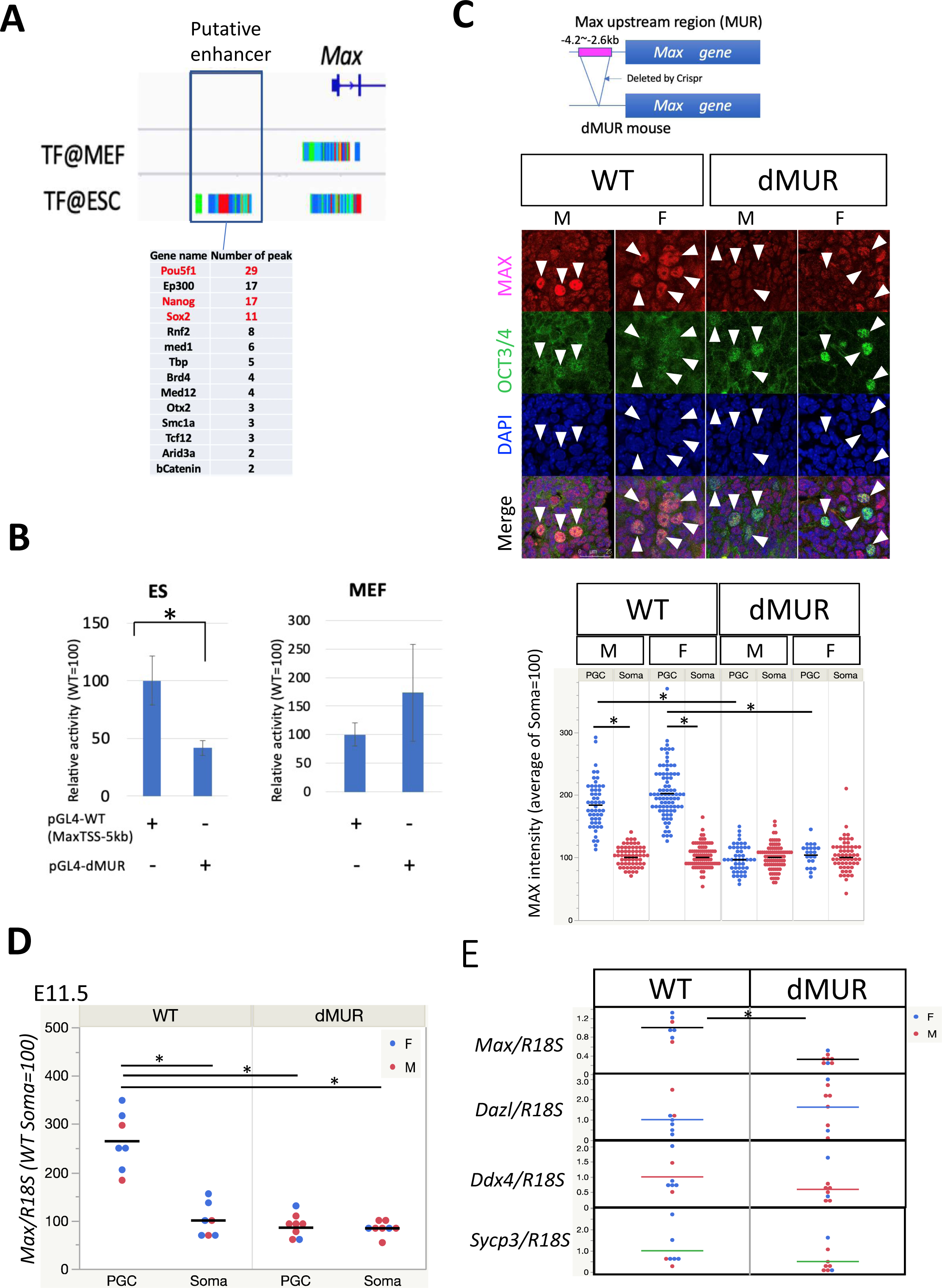
Identification of the positive regulatory region of the *Max* gene that functions commonly in ESCs and premeiotic germ cells. Computational analysis was used to search for the potential regulatory region that supports high *Max* gene expression in ESCs. Analysis of ChIP-Atlas metadata identified a genomic region that recruited representative pluripotent cell-specific transcription factors upstream of the *Max* gene. (B) Examination of transcriptional stimulating activity of MUR by reporter assays. Two luciferase reporter plasmids with or without MUR were individually introduced into ESCs and mouse embryonic fibroblasts by transfection together with an internal control plasmid carrying the *Renilla reniformis* luciferase gene. A dual-Glo luciferase assay was conducted at 48 h post-transfection. Data represent the mean ± standard deviation of three independent experiments with comparable results. Student’s t-tests were conducted. The differences in luciferase activity due to the presence of MUR was statistically significant in ESCs (*P*-value, 0.011; marked by an asterisk), but not in mouse embryonic fibroblasts (*P*-value, 0.217). (C) Immunohistochemical analyses of Max protein in germ cells of wildtype and dMUR mutant mice at E11.5. Genetic manipulation conducted in dMUR mice is illustrated in the upper panel. Double immunostaining of MAX and OCT4 in sections of genital ridges from wildtype and dMUR mice is shown. Arrowhead indicates each Oct4-positive germ cell. Lower panel shows a plot of the intensity of individual Max protein signals quantified by ImageJ software as described in Fig. 1C. The mean of the Max signal intensity in somatic cells was arbitrarily set to 100. (D) Effect of MUR deletion on *Max* mRNA levels in germ cells at E11.5. *Max* mRNA levels in germ and somatic cells of wildtype and dMUR mutant mice were quantified by quantitative PCR. Blue and red dots indicate data from female and male genital ridges, respectively. The *R18S* RNA level was used as an internal control. The asterisk indicates *P*-value for the statistical difference is less than 0.05. (E) Effect of MUR deletion on meiosis-related gene expression in germ cells at E11.5. Expression levels of *Dazl*, *Ddx4*, and *Sycp3* in germ cells of wildtype and dMUR mutant mice were quantified by quantitative PCR using the *R18S* RNA expression level as an internal control. Expression levels of *Max* and meiosis-related genes in wildtype germ cells were set to 1. No statistically significant difference was evident in expression levels between wildtype and dMUR mutant mice except for *Max* expression (*P*-value, 2.87x10^-6^; marked by an asterisk).

## Discussion

We and others have previously demonstrated that expression of meiosis-related genes is strongly repressed by PRC1.6 in ESCs with the potential to undergo at least the early stages of meiotic prophase I ectopically (Kehoe et al. 2008; Hisada et al. 2012; Qin et al. 2012; Leseva et al. 2013; Maeda et al. 2013; Suzuki et al. 2016; Yang et al. 2016; Endoh et al. 2017; Huang et al. 2018; Stielow et al. 2018; Uranishi et al. 2021). We have also reported that reducing *Max* gene expression in germline stem cells by short hairpin RNA induces meiosis-related gene expression and preleptotene/leptotene-like cytological changes (Suzuki et al. 2016). These findings imply that PRC1.6 blocks meiosis in ESCs and germ cells, but the physiological relevance is unclear. Our present study that characterized germ cells *in vivo* substantially strengthens this notion. Indeed, conditional *Max* gene knockout in PGCs induced precocious and ectopic expression of SYCP3 in female and male PGCs, respectively. However, female and male germ cells subjected to this forced initiation of meiosis failed to undergo subsequent processes progressively. Indeed, MAX depletion in female germ cells did not coordinate with physiological meiotic onset that occurs at approximately 13.5 d.p.c. to further drive meiotic progression, but meiosis was stalled mainly around the stage of transition from leptotene to zygotene. In terms of *Max*-null male germ cells, the vast majority were positive for SYCP3 before sexual differentiation, but such a high percentage of SYCP3-positive cells among *Max*-null germ cells subsequently declined substantially, and SYCP3-positive cells were hardly detected at E13.5. Our data also showed that male and female germ cells subject to forced meiotic induction, were eventually eliminated by apoptosis.

An additional important finding of this study was the dynamic changes of *Max* expression in germ cells, especially because *Max* expression level is generally known to be comparable across numerous cell types (Henriksson and Luscher 1996; Grandori et al. 2000). Indeed, female germ cells maintained approximately two-fold higher expression levels of *Max* than surrounding somatic cells in the mitotically active state, but *Max* expression declined abruptly at the timing of or immediately prior to their meiotic onset. Because this decline was assumed to be linked to an impediment of the transcription-repressing activity of PRC1.6, the decline was considered to be an important prerequisite step for germ cells to undergo meiosis. Moreover, we assume that MUR identified as a regulatory region that sustained high *Max* expression in ESCs and mitotically active germ cells substantially contributed to the subsequent downregulation of *Max* expression in germ cells, because expression of genes encoding Oct4 and other pluripotency factors, which probably conferred transcription-enhancing activity on MUR, was almost completely absent during this period. Our data also revealed that *Max* regained high expression at approximately the zygotene stage in female germ cells, which may lead to functional recovery of Max-dependent activities, i.e., Myc and PRC1.6 activities. In conjunction with the finding that *Max*-null male and female germ cells did not undergo meiotic processes progressively, but were stalled during these processes, it was assumed that recovery of Myc and/or PRC1.6 from their inactivated state via upregulation of *Max* expression was also required for germ cells to complete meiosis. Therefore, we will conduct future studies to understand how *Max* regains high expression from its low expression state during meiotic processes and how *Max* regaining high expression contributes to meiosis progression.

## Methods

### Plasmid construction and luciferase assays

The 5ʹ flanking region of *Max* was amplified by PCR from genomic DNA of wildtype mouse and mutant mouse lacking MUR using the following oligonucleotides:

5ʹ-GGTACCCCAGGAAGGATAGGCTGTGACG-3ʹ

5ʹ-GTCGACGTAGTCCTCGAGCGTCGGAT-3ʹ.

Transfection of plasmids and subsequent luciferase assays were conducted as described previously (Tomioka et al. 2002).

### Mice

Oct4 dPE-CreERT2 transgenic mice were purchased from RIKEN BRC (BRC No. RBRC06509) (Yeom et al. 1996; Yoshimizu et al. 1999). Mice with a homozygous *Max*-floxed allele (*Max*^fl/fl^ mice) (Mathsyaraja et al. 2019) were bred with Oct4 dPE-CreERT2 transgenic mice, and the resulting compound mice were bred with *Max*^fl/fl^ mice to obtain Max^fl/fl^;Oct4dPE-CreERT2^+^ mice. To generate mice lacking MUR, we used guide RNAs 5′-CCCACTGAAACATCGCCCCATGG-3′ and 5′-TCTTCACAGGCTAAGATCTCAGG-3′. Unfertilized oocytes were collected from female C57BL/6J mice (>10 weeks old) superovulated with gonadotropin and chorionic gonadotropin, and subjected to *in vitro* fertilization with sperm from C57BL/6J mice. At 5 hours post-treatment, resultant zygotes were subjected to introduction of the guide RNAs (5 ng/μl) and GeneArt Platinum Cas9 Nuclease (100 ng/μl) (Thermo Fisher Scientific, MA, USA) by electroporation using a NEPA 21 electroplater (NEPAGNENE, Chiba, Japan) as described by Sato et al. (Sato et al. 2018). Two-cell stage embryos developed by subsequent *in vitro* culture were transferred into oviducts of pseudopregnant ICR female mice for development and delivery. One of the originally obtained heterozygous dMUR mutant mice was used as a founder to maintain the colony by crossing with wildtype C57BL/6J mice. Heterozygous mutant mice were backcrossed three times before being used to generate homozygous mutant mice by heterozygous intercrosses. To distinguish between the wildtype and deletion, PCR was conducted using the following oligonucleotides.

Wildtype allele with the MUR sequence:

Forward: 5ʹ-TTTTGGCTGGACTGGCTAAC-3ʹ; reverse: 5ʹ-TTTTGGCTGGACTGGCTAAC-3ʹ

Mutant allele without the MUR sequence:

Forward: 5ʹ-TTTTGGCTGGACTGGCTAAC-3ʹ; reverse: 5’-TTTTGGCTGGACTGGCTAAC-3ʹ.

Animal experimentation was carried out in strict accordance with international and institutional guidelines. The protocol was approved by the Institutional Review Board on the Ethics of Animal Experiments of Saitama Medical University (Permission numbers: 3136, 3137 and 3138).

### Isolation of germ cells by MACS

To isolate germ cells by magnetic activated cell sorting (MACS), gonads at E10.5–14.5 were dissected from embryos and dissociated to single cells using 0.05% trypsin-EDTA in PBS after removal of the mesonephros. After treatment with MACS buffer (2 mM EDTA and 0.5% BSA in PBS), germ cells were recovered using an anti-SSEA-1 antibody conjugated to MicroBeads (Miltenyi Biotec, 130-094-530) in accordance with the manufacturer’s instructions.

### Immunostaining

For immunohistochemical analyses, genital ridges isolated from embryos (E10.5–14.5) were fixed in 4% paraformaldehyde, embedded in OCT compound, and then frozen. Sections prepared from the frozen tissues were mounted on glass slides and subjected to immunostaining as described previously (Suzuki et al. 2016). The primary antibodies used were as follows: anti-GENA (TRA98) (BioAcademia, 73-003); anti-STRA8 (Abcam, ab49602); anti-MAX (sc-197), anti-OCT4 (sc5279) from Santa Cruz Biotechnology; anti-γH2AX (Cell Signaling Technology, #2577); anti-cPARP (BD Pharmingen, 558710). Anti-GENA and STRA8 antibodies were used at a 1:400 dilution, and the anti-γH2AX antibody was used at a 1:1000 dilution. Other antibodies were used at a 1:100 dilution. Immunostained tissues/cells were observed under a confocal laser scanning microscope (TCS SP8, Leica Microsystems). The average staining intensities of MAX and SYCP3 in somatic cells were arbitrarily set to 100.

### Immunofluorescence staining for stage classification of germ cells in meiotic prophase

Preparation of nuclear spreads of germ cells was conducted in accordance with the method described by Hwang et al. (Hwang et al. 2018) with some modification. Briefly, after washing with PBS, gonads recovered from embryos were immersed in a low-tension buffer (17 mM trisodium citrate dihydrate citrate, 50 mM sucrose, 30 mM Tris-HCl, pH 8.2, 1 mM EDTA, 0.5 mM DTT, and 0.1 mM PMSF) for 7 minutes. Each pair of gonads was suspended in 50 μl drops of 100 mM sucrose on a glass slide and gently dispersed by pipetting. The solution containing gonadal cells was then dropped onto slides soaked in a fixative solution (1% PFA and 0.15% Triton X-100) and pipetted to spread the cells. After repeated cell spreading, the glass slide was incubated overnight at room temperature in a humidified box and then dried at room temperature. The cells were then applied to immunocytochemical analyses using antibodies against SYCP3 and MAX.

### Quantitative reverse transcription-PCR analysis

An RNeasy Micro Kit (QIAGEN) was used for total RNA preparation from germ cells isolated using MACS. The recovered RNAs were reverse transcribed to cDNA using an FSQ-301 kit (Toyobo). TaqMan-based quantitative PCR was performed using the StepOnePlus Real-Time PCR System (Applied Biosystems). All samples were normalized to the expression level of *bActin or RN18s*. TaqMan probes used are as follows.

*Max*, Mm00484802_g1; *Sycp3*, Mm00488519_m1; GM4969 (*Meiosin*), Mm01305445_m1; *Dazl*, Mm03053726_s1; *Ddx4*, Mm00802445_m1; Foxl2, Mm00843544_s1; Sox9, Mm02619580_g1; *bActin*, Mm02619580_g1; *RN18s*, Mm03928990_g1.

### DNA microarray and GO analyses

Biotin-labeled cDNA was synthesized and then hybridized to Affymetrix GeneChip Mouse Genome 430 2.0 arrays (Affymetrix) in accordance with the manufacturer’s instructions. Microarray expression data were background subtracted and normalized by the robust multiarray analysis method using statistical software R. GO analyses were conducted using DAVID web tools (http://david.abcc.ncifcrf.gov). The selected GO terms were subjected to further analyses using AmiGO1 (http://amigo1.geneontology.org/cgi-bin/amigo/go.cgi) and REVIGO (http://revigo.irb.hr) websites to eliminate synonymous terms.

### Accession number

DNA microarray data have been deposited in the NCBI Gene Expression Omnibus under accession number GSE233556.

## Acknowledgments

We are indebted to Ms. Pei Feng Cheng (Fred Hutchinson Cancer Research Center) and Dr. Atsushi Suzuki (Yokohama National University) for mice with targeted *Max* deletion and Oct4dPE-CreERT2 mice, respectively. We are also grateful to Kazumi Ubukata for his technical assistance and Mitchell Arico from Edanz (https://jp.edanz.com/ac) for editing a draft of this manuscript. This study was supported in part by grants from the Japan Society for the Promotion of Science (JSPS) KAKENHI (Grant numbers: 21K06843, 23K06391, and 23H02678) to AS, KU, and AO, respectively, and a grant from National Institute of Health (Grant number: NIH/NCI R35 CA231989) to RNE. This study was also supported in part by grants from the Takeda Science Foundation to AS and KU and by internal grants from Saitama Medical University (Grant numbers: 20-G-1-01 and 21-B-1-16) to AS and KU, respectively.

## Author contributions

A.S., Conception and design, conduct of experiments, data analysis, manuscript writing; K.U., M.N., and Y.M., Conduct of experiments and data analysis; S.M., and S.T., Genetic manipulation of mice; R.E., Conception and design, and manuscript writing, A.O., Conception and design, financial support, manuscript writing, and final approval of the manuscript.

