## Supplemental Figures for "Ablation of *Max* expression induces meiotic onset in sexually undifferentiated germ cells"

A

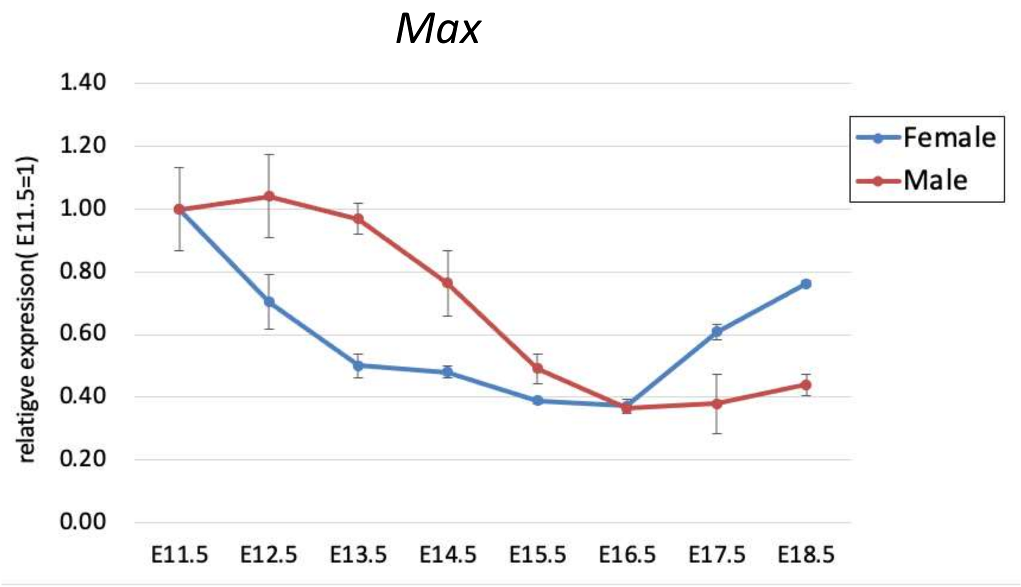


B

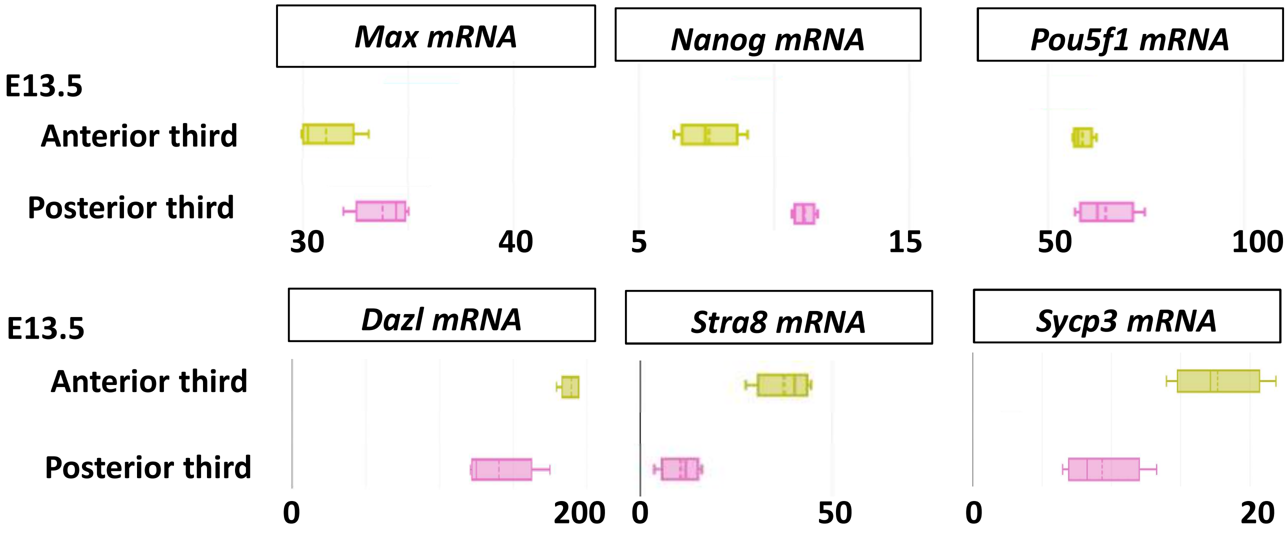


Supplemental figure. S1. Spatiotemporal alteration of expression profiles of *Max*, pluripotency and meiosis-related genes in germ cells (related to Figure 1)

(A)Expression dynamics of *Max* in male and female germ cells during mid- and late-embryonic stages. Data were extracted from the published microarray data (GSE23322) (Sabour et al. 2011). *Max* expression level in male and female germ cells at E11.5 was respectively set to 1.

(B) Expression levels of *Max*, *Nanog*, *Oct4*, *Dazl*, *Stra8* and *Sycp3* genes at anterior and posterior portions of female gonads at E13.5. Box plots were generated using data reported by Soh et al. (2015) via the website of The REPO GENOMICS VIEWER (https://rgv.genouest.org/).

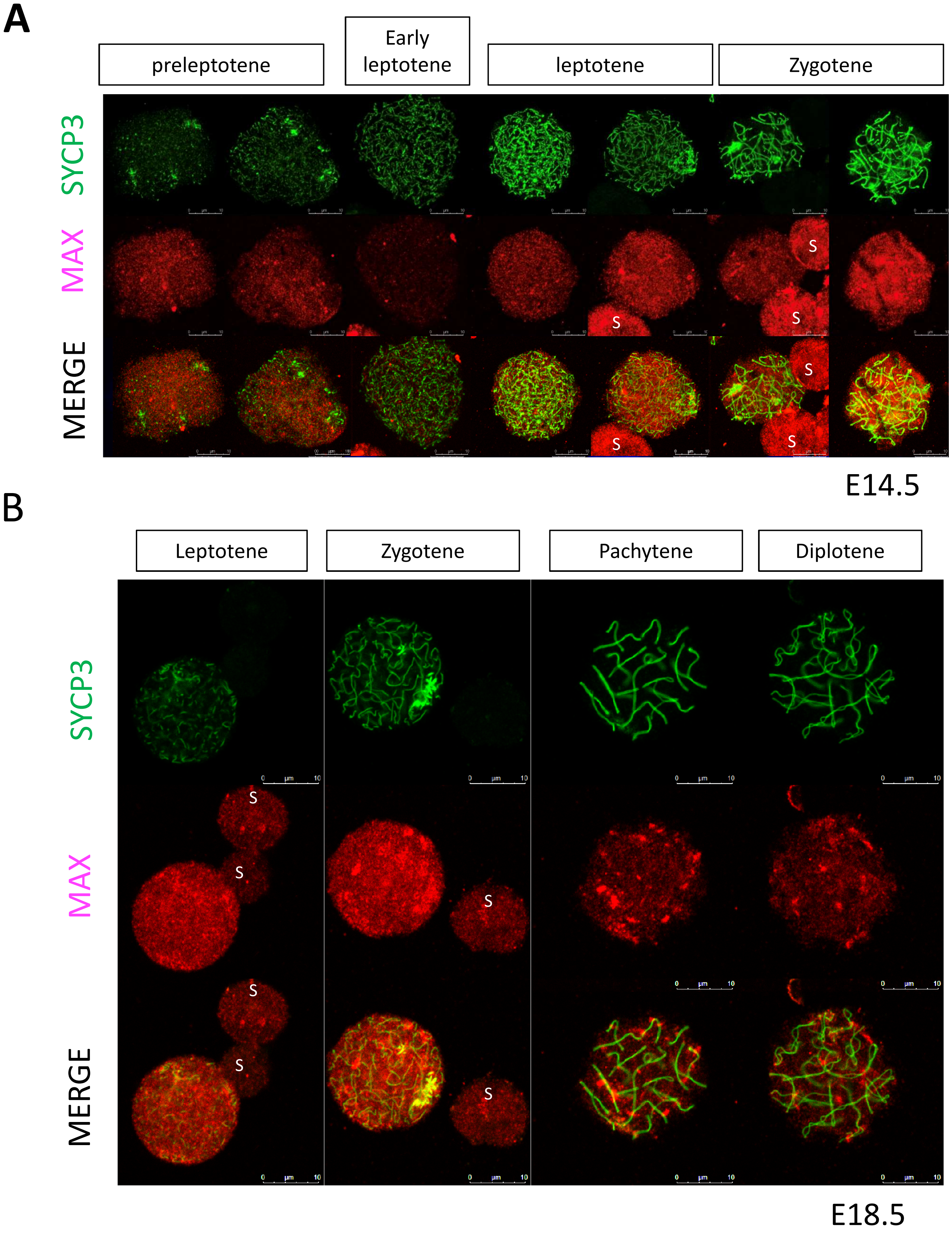


Supplemental figure 2. Immuno-cytochemical analyses of MAX and SYCP3 with female germ cells during their meiotic prophase I (related to Figure 1)

(A-B) Nuclear spreads of female germ cells at E14.5 (A) and 18.5 (B) were subjected to immuno-cytochemical analyses using anti-SYCP3 and MAX antibodies. Stage of each germ cell was determined according to the SYCP3 staining pattern. S: somatic cells

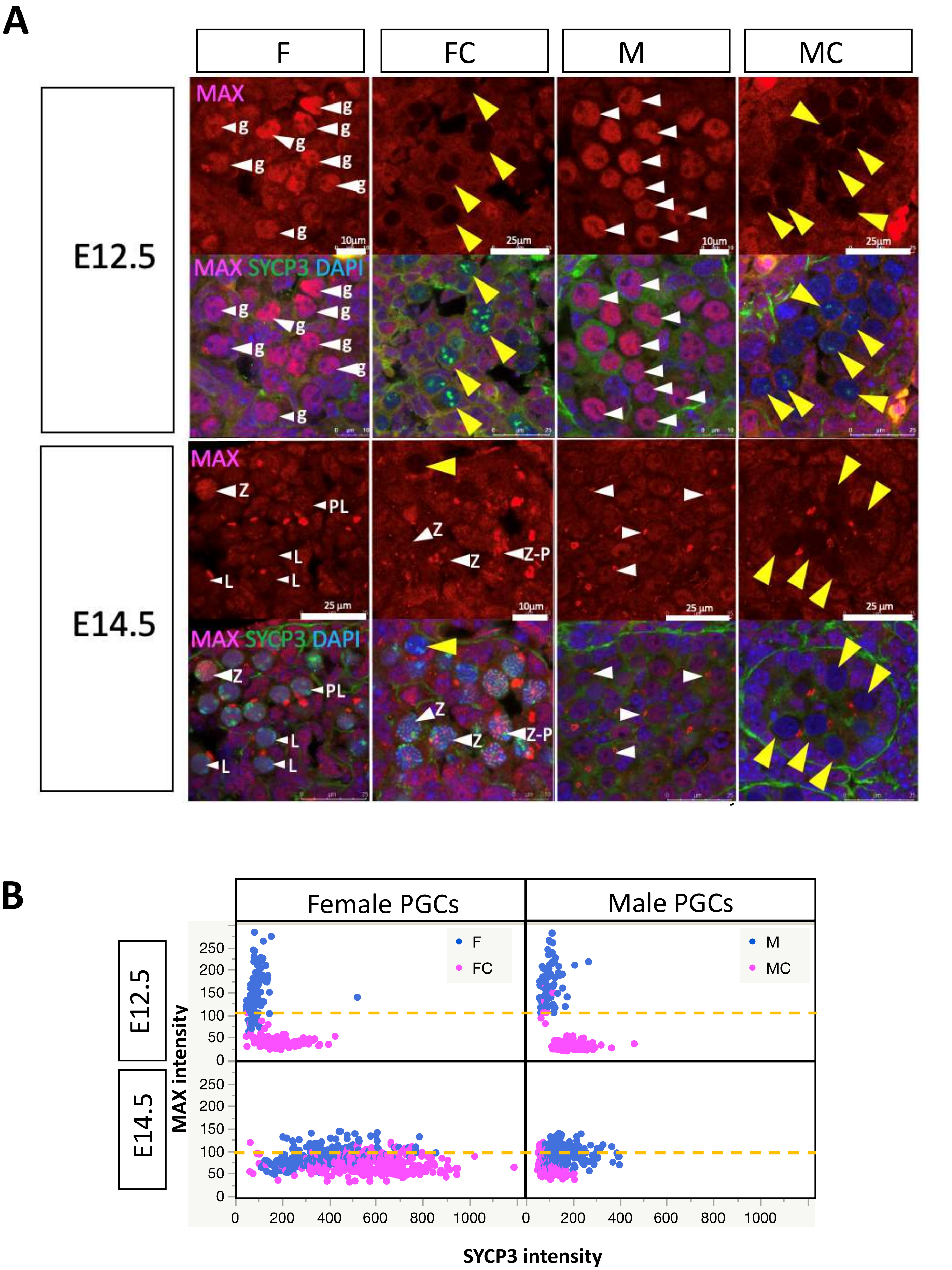
A
F
FC
M
MC
E12.5
E14.5
MAX
MAX SYCP3 DAPI
g
Z
PL
L
Z-P
10μm
25μm
25 μm
B
Female PGCs
Male PGCs
F
FC
M
MC
MAX intensity
SYCP3 intensity
0
50
100
150
200
250
200
400
600
800
1000

Supplemental Figure 3. Immuno-histochemical analyses of germ cells subjected to Cre-mediated *Max* gene disruption and their control counterparts for MAX and SYCP3 (related to Figure 2).
(A)Double immunostaining analysis. Embryonic female and male germ cells at E12.5 and E14.5 that were either subjected to Cre-mediated *Max* gene disruption by tamoxifen administration or control counterparts carrying no CreERT2 cDNA were immune-stained for MAX (Magenta) and SYCP3 (Green) and counterstained with DAPI (Blue). Each germ cell is pointed by an arrowhead. Yellow arrowhead is used to point germ cells whose Max protein signal is completely null because of tamoxifen administration-mediated *Max* gene disruption. g, PL, L and Z indicate undifferentiated germ cells, preleptotene, leptotene and zygotene, respectively.
(B) Scatter plot representing intensity of Max and Sycp3 staining signals in individual male and female germ cells at E12.5 and 14.5 that were either subjected to tamoxifen treatment for *Max* gene disruption (MC and FC) or control counterparts carrying no CreERT2 cDNA (M and F).

**A**

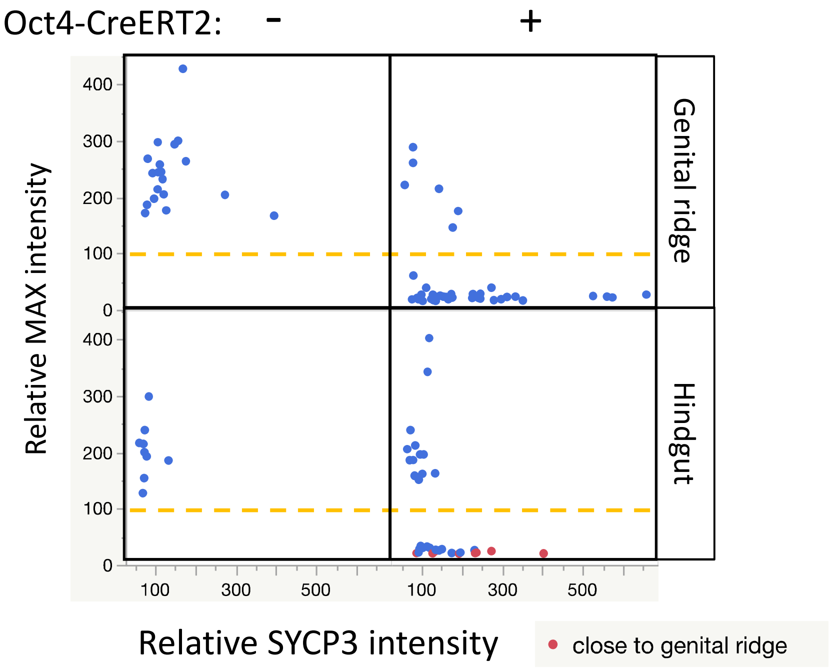


**B**

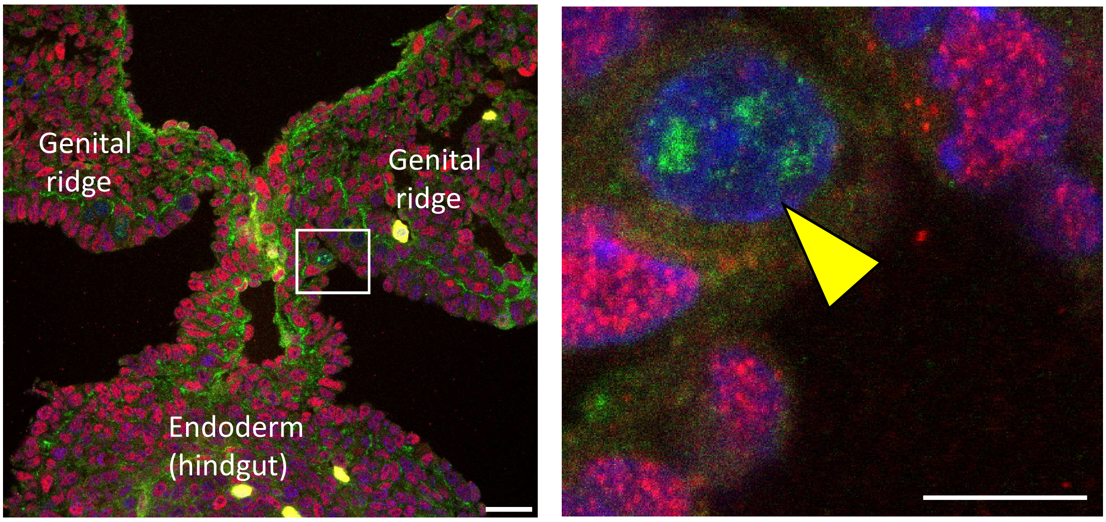


Supplemental Figure 4. Effect of *Max* expression ablation in the migrating PGCs at E10.5 (related to Figure 2)

(A)Scatter plot of intensity of Max and Sycp3 staining signals in individual male and female germ cells at E10.5 in genital ridge and hindgut that were either subjected to *Max* gene disruption by tamoxifen administration at E8.5 or control counterparts carrying no CreERT2 cDNA. Protein signal intensity was determined using ImageJ software as in Figure 1C. The intensity of Max and Sycp3 signals obtained with somatic cells were respectively set to 100. Data from germ cells that were located within hindgut, but close to genital ridge were marked with pink dot.

(B) Representative example of SYCP3-staining positive migrating germ cells that had migrated in the vicinity of genital ridge. A region containing SYCP3-positive migrating germ cells is marked with a rectangle and its magnified image is shown in the right. Each scale bar corresponds to 10 mm.

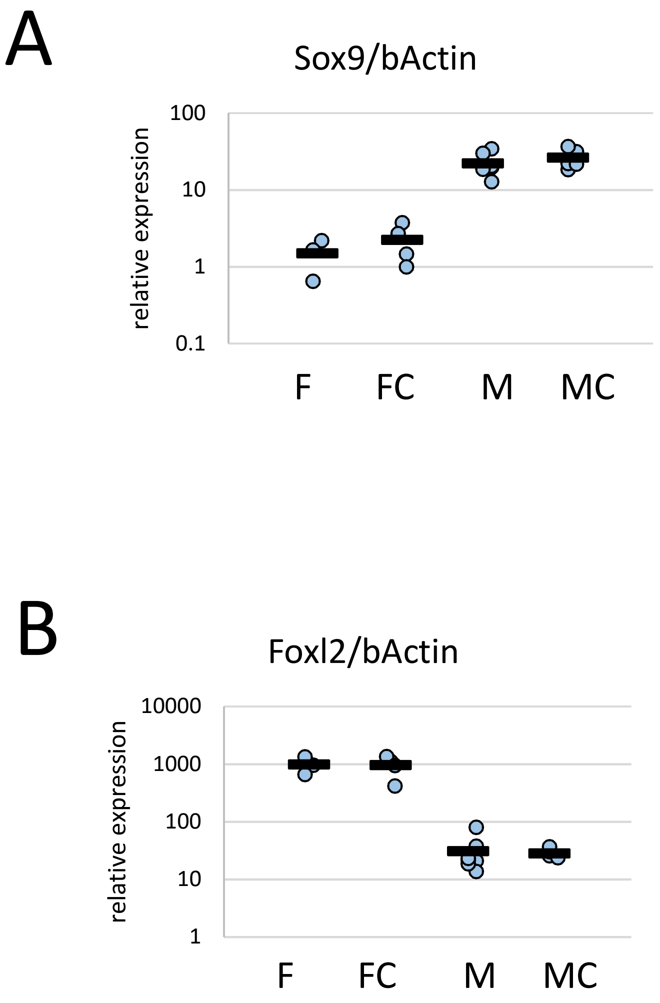


Supplemental Figure 5. *Max* gene disruption was not accompanied with alteration in magnitude of sex differentiation-related gene expression level (related to Figure 3)
(A,B) Expression levels of *Sox9* (A) and *Foxl2* (B) genes were quantified with RNAs from whole gonads of E12.5 embryos that were subjected to Cre-mediated *Max* gene disruption by tamoxifen administration at E8.5 (FC and MC) or control counterparts carrying no CreERT2 cDNAs (F and M).

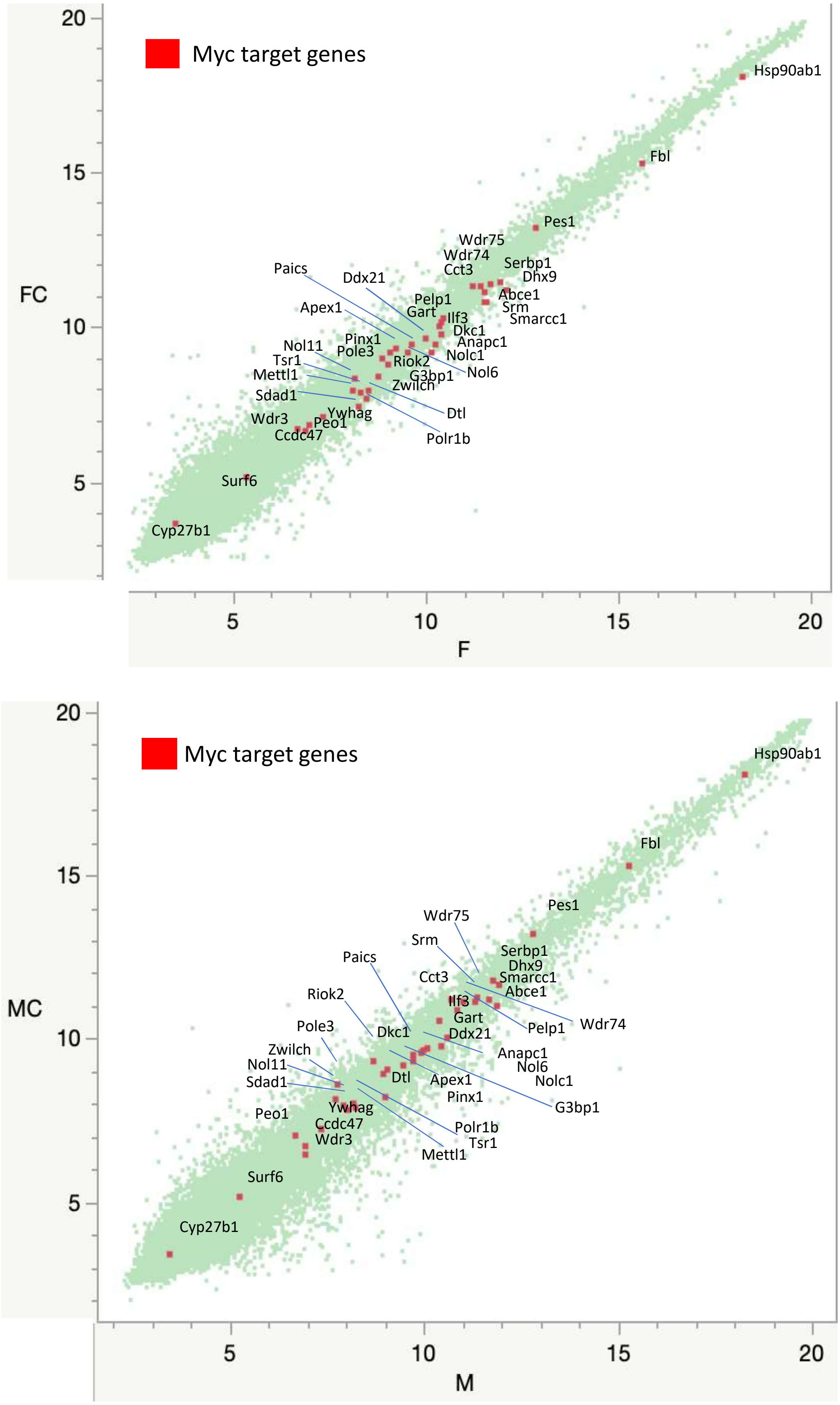


Supplemental Figure 6. Effect of *Max* expression ablation on Myc target genes (related to Figure 3)
Data of genes designated as members of cell-type independent core Myc targets (Ji et al. 2011) were marked with red dots with gene symbols on scatter plots of DNA microarray data shown in Figure 3A. Upper and lower panels show data obtained with female and male germ cells, respectively.

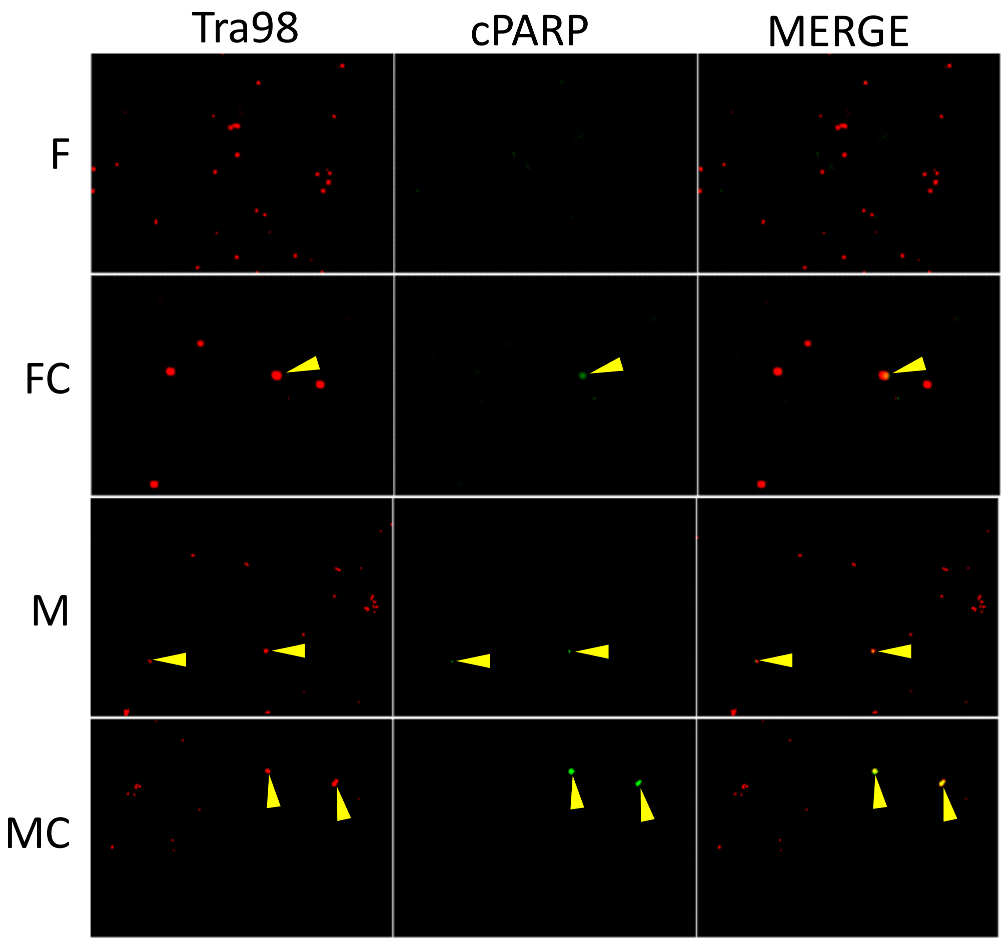


Supplemental Figure 7. Germ cells that onset meiosis artificially by the force *Max* expression ablation showed apoptotic phenotype (related to Figure 4).
Representative images of double immuno-stained TRA98 and cPARP of germ cells from whole gonads of embryos that were subjected to Cre-mediated *Max* gene disruption by tamoxifen administration at E8.5 (FC and MC) or control counterparts carrying no CreERT2 cDNA (F and M). TRA98/cPARP double-positive germ cells were indicated with yellow arrowheads.

**A**

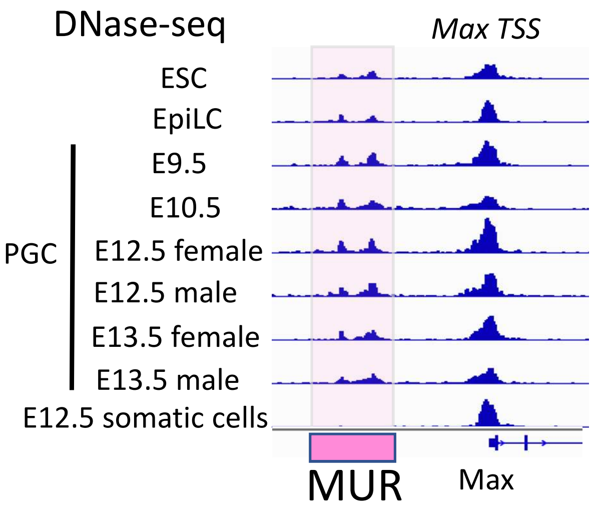


**B**

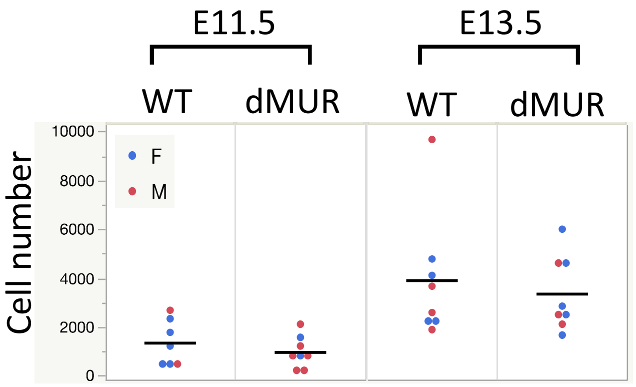


Supplemental Figure 8. DNase hypersensitivity of MUR in pluripotent cells and germ cells at pre-meiotic stage and the consequence of its loss on total germ cell number (related to Figure 5).

(A) DNase hypersensitivity of the region around transcription start site of *Max* gene and its 5'-flanking region. Publicly available data for ESC (SRX3606939), EpiLC (SRX3606943) and PGCs at E9.5 (SRX3606915), E10.5 (SRX3606917), E12.5 female (SRX3606919), E12.5 male (SRX3606923), E13.5 female (SRX3606925), E13.5 male (SRX3606927), and somatic cells at E12.5 (SRX3606921) reported by Li et al. (2018) were used to generate this snapshot.

(B) Total germ cell numbers that were assessed by TRA98 staining in gonads of wild-type and mutant mice that lack MUR at E11.5 and 13.5. No statistical difference in total germ cell number between wild-type and dMUR mutant mice was evident.
